# Population-level genome sequencing reveals distinct Mycobacterium tuberculosis intrahost mutational trajectories in simian immunodeficiency virus co-infected and antiretroviral treated non-human primates

**DOI:** 10.64898/2026.04.03.714442

**Authors:** Michael C. Chao, Michael R. Chase, Shoko Wakabayashi, Andrew Vickers, Byron Roman, Forrest Hopkins, Peter H. Culviner, Maximillan G. Marin, Pauline Maiello, Collin R. Diedrich, Zandrea Ambrose, Philana Ling Lin, Qingyun Liu, Sarah M. Fortune

## Abstract

Whole genome sequencing of *Mycobacterium tuberculosis* (Mtb) populations from clinical samples has increasingly identified genes undergoing selection within and between hosts that drive differential infection and treatment outcomes. However, the intrahost Mtb mutational landscape—especially in the context of human immunodeficiency virus coinfection and antiretroviral therapy (ART)—remains less clear, as do the potential impacts of such mutations on Mtb infection dynamics. Here, we performed whole genome sequencing of Mtb populations isolated from 477 infected tissues across 20 non-human primates, including animals co-infected with simian immunodeficiency virus (SIV) with or without virological suppression by ART. We identified 116 mutations that emerged during infection, including those that are overrepresented within individual tissues and a subset that are shared across tissues during Mtb dissemination. We further find differential mutation trajectories across treatment groups, with higher mutation rate and bacterial outgrowth in SIV-infected hosts and increased prevalence of oxidative damage-associated mutations in coinfected animals on ART. Finally, we demonstrate a common pattern of mutation in Mtb lipid metabolism and polyketide synthase genes and identify a subset of NHP-derived mutations that have also independently arisen in human clinical isolates. Together, our population-based sequencing uncovers Mtb diversification during early infection, captures discrete bacterial dissemination events and infers differential immune pressures faced by Mtb in the setting of SIV-Mtb coinfection and ART suppression.

**Importance:** Tuberculosis remains a leading cause of death worldwide, especially in people living with HIV. How HIV infection and antiretroviral therapy impacts *Mycobacterium tuberculosis* (Mtb) intrahost evolution remains unclear. Using whole genome sequencing from hundreds of infected tissues from non-human primates, we find that simian immunodeficiency virus co-infected hosts and those receiving antiretroviral therapy exact different immune pressures on Mtb leading to different mutation rates and types of DNA damage that are incurred. Mtb mutations were enriched in genes involved in lipid metabolisms and some of these are also seen in human TB strains. This work highlights the role of immune pressure to alter bacterial pathways that may enable Mtb adaptation to the host.

## Background

Tuberculosis (TB) remains the top single infectious cause of mortality worldwide, with people living with HIV (PLHIV) bearing an outsized cost of the ongoing epidemic^1^. Studies have shown significant improvement in preventing TB mortality in this population through the widespread use of antiretroviral therapy (ART)—especially when started early—but individuals on long-term ART still retain an elevated risk of developing of active TB disease despite virologic control and stable CD4 counts^2–5^. Though studies have identified dysfunctional adaptive and innate immune responses associated with HIV/Mtb coinfection (e.g., reviewed in ^6^), the mechanisms accounting for increased TB disease progression in PLHIV remain incompletely understood. To model early *Mycobacterium tuberculosis* (Mtb) infection outcomes related to HIV coinfection, a recent study in non-human primates (NHP) established chronic simian immunodeficiency virus (SIV) infection first—including a subset of animals with virologic suppression using ART—before challenging animals with a genetically barcoded isogenic library of Mtb^7^. This work found that SIV-positive animals developed more severe TB disease, including higher bacterial burden and greater inflammation; and that early ART during SIV infection could significantly reduce these effects. However, PET-CT imaging, pathology scoring and sequencing of barcoded Mtb strains also showed that ART did not restore Mtb dissemination to levels observed in non-SIV infected controls—especially to extrapulmonary sites—suggesting that ART animals experienced continuing defects in certain aspects of early Mtb control^7^. However, the impact of SIV and ART on the bacterial intrahost evolutionary landscape in this study remained unclear.

Whole genome sequencing of Mtb clinical strains is increasingly identifying genes and pathways undergoing selection in hosts that drive differential infection and treatment outcomes^8–10^. These mutational signatures have been powerful tools for studying long term co-evolution of Mtb with human hosts^10^ and identification of novel mutations that contribute to the emergence of antibiotic resistance^11,12^, many of which have been translated into diagnostics for detection of multi- and extremely-drug resistant infections. Further, sequencing of intrahost Mtb populations can identify unfixed antibiotic-resistance mutations that emerge during treatment^13–15^ and can serve as a record of host immune pressure acting on Mtb^16,17^. However, sequencing is often limited to anatomically accessible sites such the airway, with studies suggesting there is greater Mtb diversity present in the human lung and extrapulmonary tissues compared to sputum^18,19^.

In this study, we hypothesized that sequencing Mtb genomes from anatomically diverse tissues in NHPs would comprehensively identify intrahost mutations, uncover distinct immune pressures in the context of SIV coinfection with and without ART and also highlight biologic processes that may be important for Mtb adaptation to the host milieu. Here, we performed whole genome sequencing of Mtb populations isolated from 477 infected tissues from 20 non-human primates, finding 116 Mtb mutations that arose within the host. These mutations show that the Mtb mutation rate is significantly increased in SIV-infected NHPs, while the proportion of oxidative damage-associated mutations is increased in animals on ART compared to non-SIV controls. We further show that Mtb lipid metabolism and biosynthetic genes are enriched for intrahost mutations, which we propose reflect opportunities for Mtb adaption to host nutrient conditions.

## Results

### Whole genome sequencing defines the intrahost Mtb mutational landscape

In this work, we curated a set of Mtb genomic DNA samples isolated from a prior non-human primate (NHP) study^7^, which included naïve or chronically SIV-infected animals (with or without ART) that were challenged with a barcoded library of Mtb. In prior work, the viable Mtb populations from all infected tissues (including lung, thoracic lymph node and extrapulmonary sites) were plated for genomic DNA extraction and amplicon-based sequencing was used to define the barcoded Mtb strains that were infecting each tissue^7^. In this study, we performed whole genome sequencing of this DNA from a subset of 477 intrahost Mtb populations isolated from 20 NHPs (Supplementary Table 1), including 7 that were solely infected with Mtb (TB); 4 chronically infected with SIV prior to Mtb challenge (SIV+TB); and 9 SIV-infected but virally suppressed with ART (SIV+ART+TB) only 3 days after infection (Figure 1A).

**Figure 1.**
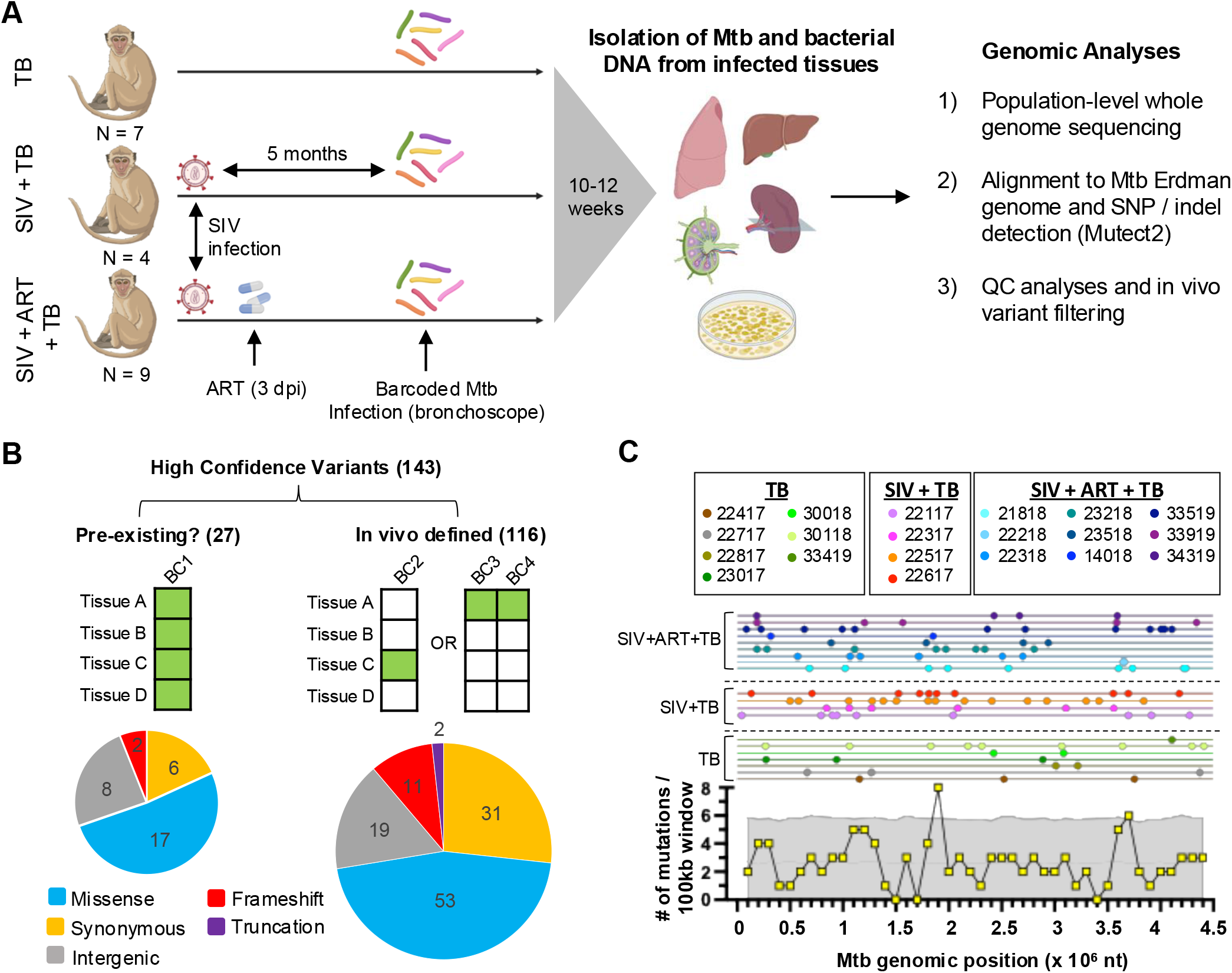
Whole genome sequencing identifies intrahost mutations. (A) Study design in which naïve, SIV infected NHPs (with and without suppressive ART initiated at 3 days post infection) were challenged with a barcoded Mtb library. All bacteria were isolated from infected tissues, genomic DNA extracted for whole genome sequencing and intrahost mutations identified. (B) Top: criteria for defining in vivo derived mutations, where every row (Tissues A-D) represents sites infected by a barcoded Mtb strain (columns BC1-4). Tissues containing a genomic mutation are colored green while white indicates presence of wild type Mtb. Bottom: Mutations are grouped by their predicted impact on protein coding. (C) Top: for each animal (lines), the genomic positions with detected mutations in any tissue are plotted (dots). Bottom: the numbers of mutations (from all NHPs) observed within 100 kb genomic windows are plotted (yellow). Grey shading indicates 2 standard deviations of the number of expected mutations in a given 100 kb window when sampling 116 random sites in the Mtb genome across 1000 simulations. The simulation was conducted after masking the repetitive sites in the genome (as was done for intrahost mutation detection^20^) but did not otherwise consider other genomic features like nucleotide identity.

Mtb sequences were mapped to a recently generated Mtb Erdman reference genome^20^, which was derived from a representative colony from the library used to infect these NHPs. Using the haplotype mutation calling tool Mutect2, we defined over 100,000 single nucleotide polymorphisms (SNPs) and small insertion and deletions (indels) across all samples, with the vast majority at low frequency and likely representing sequencing artifacts (Supplementary Figure 1A). Subsequent filtering of low coverage sites and considering only variants at >10% prevalence within tissues (see Methods) yielded 143 unique, high-confidence candidate mutations that were not present in our reference genome (Supplementary Table 2). As some variants could have arisen during library construction in vitro (e.g., see ‘BC1’ example in Figure 1B), we further compared the tissues that contained a genomic variant against previously generated barcoding data from the same samples to down select 116 mutations that were likely in vivo-derived (see Methods), as they are only found in a subset of sites infected by the same barcoded Mtb strain (e.g., BC2, BC3 and BC4 in Figure 1B). This left 27 mutations that we cannot rule out are pre-existing in the infecting library (Supplementary Table 2), so we elected to exclude these from downstream analyses.

Of the intrahost mutations, 57% were predicted to change protein coding (53 missense mutations, 11 frameshift indels and 2 in-frame internal deletions), 31 were predicted to be synonymous and 19 mutations were located within intergenic regions (Figure 1B). Cumulatively, intrahost mutations occurred across the Mtb genome, but certain genomic regions harbored more mutations than would be expected given completely random sampling (Figure 1C). Specifically, we observed local mutational maxima between nucleotides 1.8–1.9 Mb and 3.6–3.7 Mb, with the highest mutational density found between 1.8–1.9 Mb, a region that includes several polyketide synthase genes (*pks7, pks8* and *pks11*).

### SIV-infected and antiretroviral treated NHPs harbor Mtb with divergent mutational trajectories

We next calculated relative Mtb mutation frequency across treatment groups using the total number of unique mutations identified in each animal (see Methods). First, we elected to use a fixed generation time for our calculations in order to define a relative mutation burden per infection day across conditions, which aimed to understand whether the mutational frequencies were different across conditions. In Figure 2A, we found that SIV-coinfected NHPs (but not those on ART) had significantly higher mutational frequencies per day compared to Mtb only animals. Next, we considered what may be possible drivers of higher relative mutation frequency, which includes increased Mtb replication rates. In SIV-infected animals, there was a statistical trend (p=0.09) of increased Mtb replication over the course of infection (Figure 2B), as evaluated by quantifying chromosomal equivalents (CEQ)—the number of Mtb genomes in tissue homogenates representing the total live and dead bacteria burden over the course of infection. Further, there is significantly more live Mtb (CFU) in SIV coinfected animals at necropsy (Figure 2C), suggesting decreased capacity to clear Mtb.

**Figure 2.**
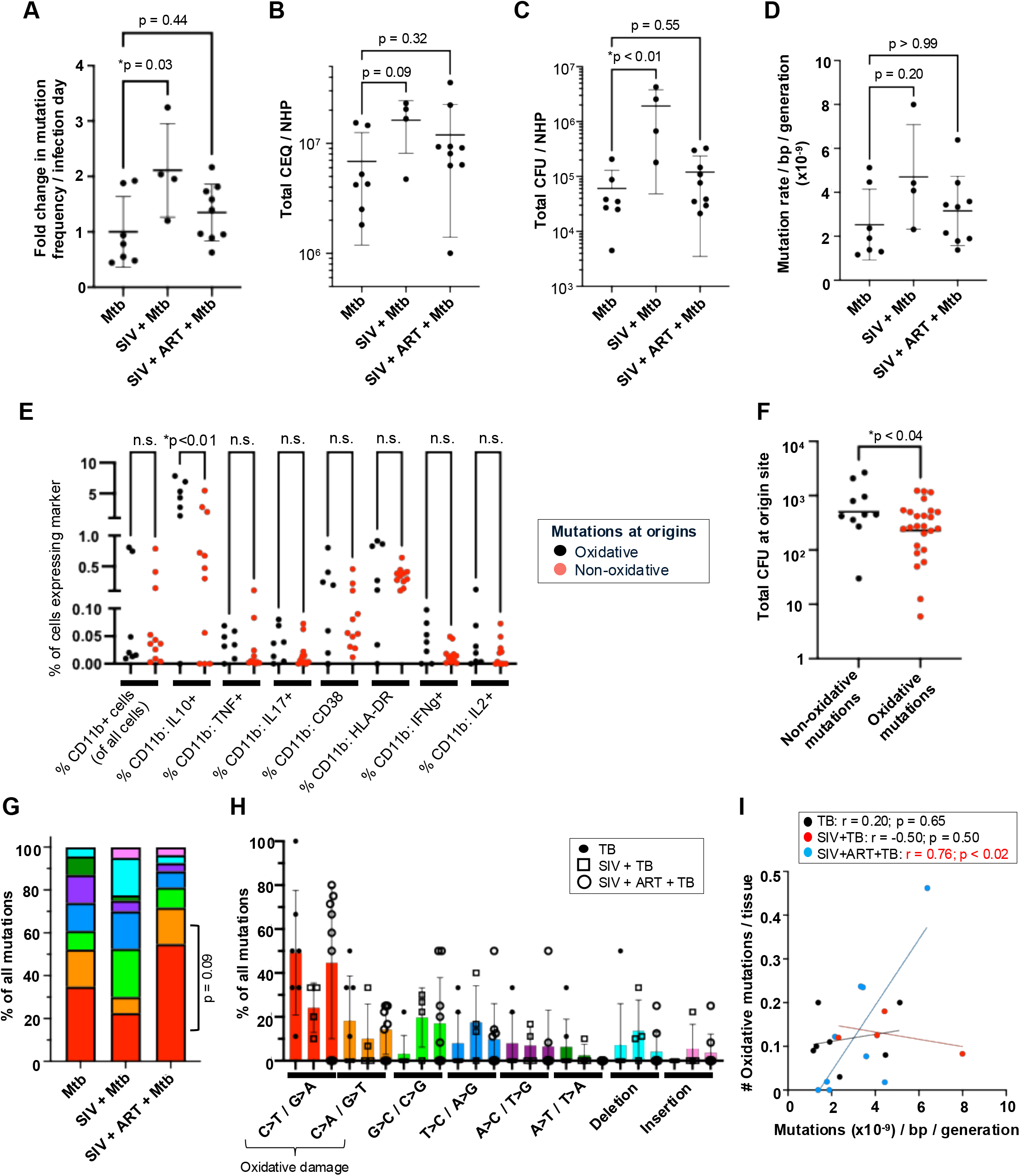
Mutational rates and profiles across SIV coinfection and ART. (A) Relative mutational rates were calculated for each animal per unit of infection time. (B) The total chromosomal equivalents (CEQs) from all infected tissues in each NHP was compared across treatment groups. (C) The total colony forming units (CFU) in each NHP at necropsy was compared across groups. (D) Mtb mutational rates per generation were calculated using an animal-specific generation time. (A-D) Statistical testing was performed using a Kruskal Wallis test with Dunn’s correction. (E) Flow cytometry profiling of myeloid cells from origin lesions containing oxidative (C>T/G>A and C>A/G>T) and non-oxidative damage mutations (all other SNPs). Statistical comparisons were conducted using One-way ANOVA with Bonferroni correction. (F) The total Mtb CFUs at necropsy for origin sites from Mtb only and Mtb + ART + SIV NHPs containing an oxidative- or non-oxidative damage mutation were compared using an unpaired t-test. (G) The percent of all mutations observed in a treatment group attributed to a distinct type of genomic mutation. P-value was defined by a Fisher’s Exact test of oxidative mutations versus non-oxidative mutations in the ART group compared to the TB only group. (H) The percent of all mutations in each NHP (dot) separated by type of mutation. (I) Correlation between the number of oxidative damage-associated mutations per NHP (normalized by number of tissues sampled) with each animal’s Mtb calculated mutation rate per generation (one-tailed Pearson correlation).

To estimate the overall Mtb replication dynamics in the outbred NHP model, and thereby evaluate the contribution of increased Mtb replication in driving Mtb diversification in SIV-coinfected NHPs, we first sought to determine the overall Mtb population doublings in each NHP by using the total unique barcodes and CEQs obtained across tissues in each animal as the starting and final Mtb population sizes, respectively (see Methods). These calculations found an average Mtb mutation rate of 2.5 x 10^-9^/bp/generation in the TB only group, which is approximately 10-fold higher than the Mtb in vitro mutation rate that was previously calculated using fluctuation tests^16^ and may reflect the contribution of host drivers of Mtb diversification. Next, we compared the intrahost replication dynamics across treatment conditions, finding that SIV-coinfected hosts support greater Mtb replication (Supplementary Figure 1C) and that Mtb have a significantly shorter generation time in SIV-coinfected hosts (Supplementary Figure 1D). When we now account for differences in generation time, we do not observe a significant difference in mutation rate per generation in SIV co-infected hosts (Figure 2D), which suggests that the elevated number of observed mutations in SIV-coinfected animals may be mainly driven by enhanced bacterial replication capacity in these immunocompromised hosts.

We next evaluated DNA damage due to host immune attack as another driver of intrahost mutation, as prior work suggested that the majority of intrahost Mtb mutations in latently infected NHPs and in sputum of active TB patients are mutations associated oxidative damage^16,17^—namely, C>T/G>A transitions and A>G/C>A transversions reflecting cytosine deamination^21^ and 8-oxo-guanine formation^22^, respectively. However, oxidative damage can also emerge in vitro during genomic DNA extraction^23^, so we explored whether tissues harboring oxidative mutations are associated with markers of immune activation or ability to control Mtb. To test these hypotheses, we first down-selected 54 ‘origin’ tissues (Figure 3, boxed nodes) where genomic mutations likely arose (see Methods), as mutations found in some tissues could have hitchhiked through Mtb dissemination. Next, we found that 18 of these lesions (11 containing oxidative mutations and 7 containing non-oxidative mutations) had been previously profiled by flow cytometry^7^, so we assessed whether myeloid cells had differential expression of cytokines in oxidative and non-oxidative damage associated lesions. In Figure 2E, we find that oxidative mutation harboring lesions had significantly lower expression of IL-10, which is an immunomodulatory cytokine that has been shown to impair macrophage immune activation and Mtb control^24,25^. This may suggest that non-oxidative lesions have dampened capacity to control Mtb; and when we compared the Mtb burden at origin sites, we find that sites harboring oxidative mutations have significantly lower CFUs compared to sites associated with non-oxidative mutations (Figure 2F). Thus, while it is difficult to causally link oxidative-damage associated mutations to immune attack without further interventional studies, our data are consistent with Mtb oxidative mutations arising in tissues associated with enhanced immune response.

**Figure 3.**
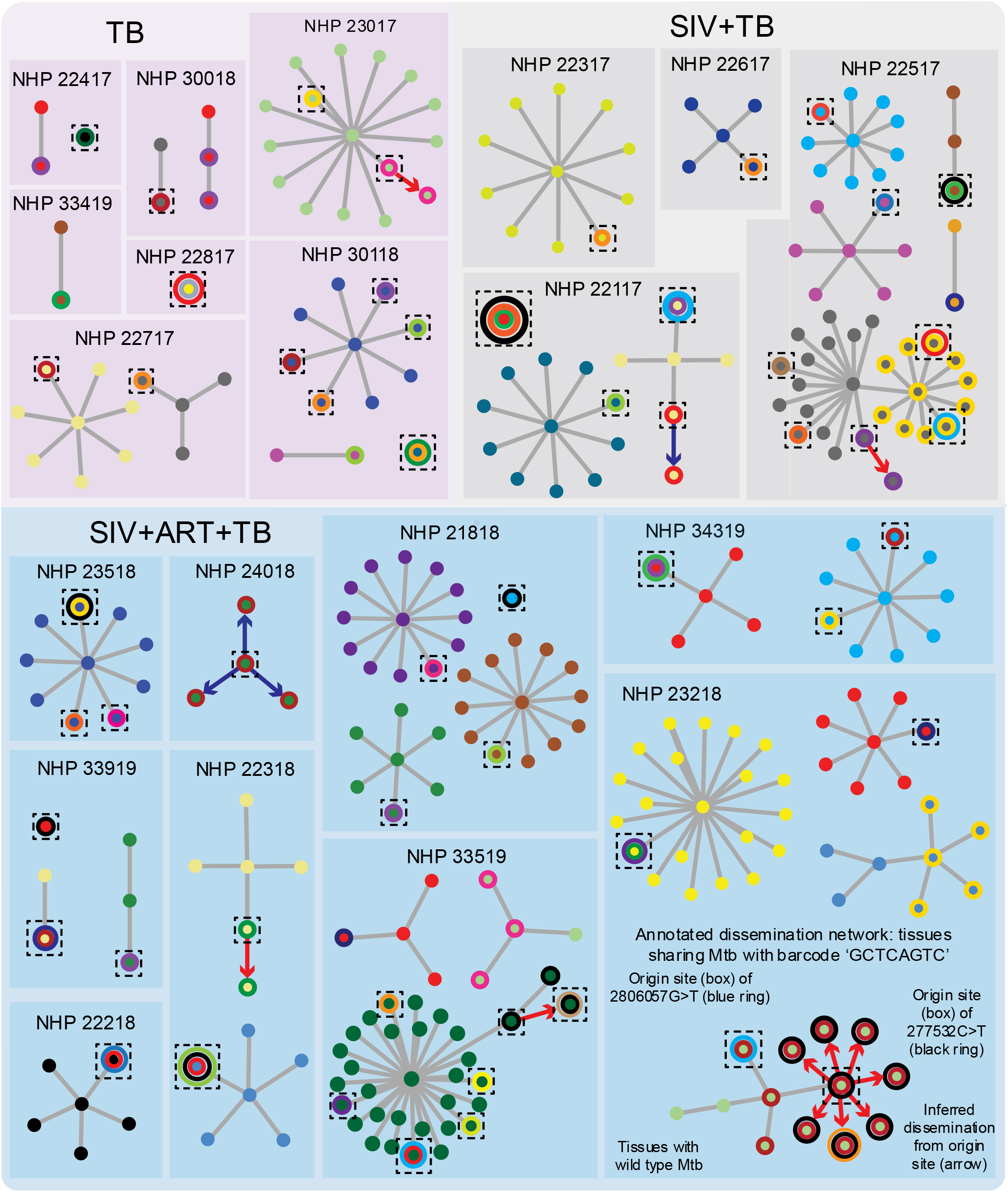
In vivo variants uncover discrete dissemination events within barcoded Mtb dissemination networks. Colored nodes denote tissues sharing the same Mtb strain as defined from prior barcode sequencing in ref^7^. Rings indicate Mtb mutations present in that tissue, with each color representing a different genomic mutation. Tissues with one barcoded Mtb strain and a mixture of wildtype and mutant alleles likely serve as ‘origin’ sites, which are denoted as boxes. Inferred dissemination events from origin sites are indicated by arrows (red arrows indicate origin site is a lung tissue disseminating to other lung tissues, while blue arrows indicate origin site is a thoracic lymph node disseminating to lung tissues).

We next compared the prevalence of oxidative mutations across treatment conditions, finding that SIV+ART+TB (but not SIV+TB) animals trended (p = 0.09) towards more oxidative mutations compared to Mtb only animals (Figure 2G). This signal was driven by a majority of ART animals harboring elevated frequencies of C>T/G>A and C>A/G>T mutations (Figure 2F). Further, there is significant correlation between the prevalence of oxidative damage-associated mutations with in vivo Mtb mutation rates in SIV+ART+TB animals, but not for the TB only or SIV+TB animals (Figure 2G). One hypothesis could be that Mtb is experiencing a range of oxidative damage in the SIV+ART+TB animals and this can lead to increased Mtb mutation rates in a subset of animals. If this oxidative damage is host-directed, this may be recorded as earlier appearance of mutations at origin sites; and consistent with this, we find that lung granulomas with intrahost mutations were detected significantly earlier by PET-CT in the SIV+ART+TB animals compared to the TB only group (Supplementary Figure 1E). Together, the data suggest that despite ART viral suppression leading to similar bacterial outcomes (Figure 2A-C), the Mtb intrahost mutational trajectory in these animals may be distinct from non-SIV animals, instead favoring a pattern of increased oxidative-associated damage.

### Intrahost mutations uncover dissemination events within barcoded Mtb dissemination networks

By integrating our whole genome sequencing with previous barcoding data, we were able to link 79 intrahost mutations with the Mtb barcoded strain that harbors these variants (see Methods). Of these, 67 mutations were found in a single tissue (considering all sampled lung, lymph node and extrapulmonary sites) while 13 variants were shared across tissues (Figure 3), suggesting they likely arose once and were spread through subsequent dissemination events. As only some tissues colonized by the same barcoded Mtb strain may harbor a given intrahost mutation, we used in vivo variants as additional ‘barcodes’ with which to infer the order of dissemination events in each Mtb dissemination network. As an annotated example, we identified 4 in vivo variants occurring in the same Mtb barcoded strain (Q25_CGTCAGTC) background in NHP 23218 (Figure 3, bottom right). By mapping the intrahost variants present and absent in these tissues, we are able to infer at least 3 distinct dissemination events: first, two tissues represent dissemination of the wildtype parent strain; second, a TCCCAC>T deletion at position 182022 (red ring) emerged and was then disseminated to 12 tissues; and third, a new 277532_C>T variant (black ring) arose on top of the TCCCAC>T deletion and both mutations were then disseminated to 7 other sites. Finally, two variants—one (2554553_C>T, orange ring) that appeared in an otherwise wildtype background and one (2806057_G>T, blue ring) that appeared on top of the TCCCAC>T deletion—emerged independently in single tissues likely after these prior dissemination events.

Similar branched networks were inferable across other animals (e.g., NHP 22517 (grey), NHP 33519 (green), NHP 24018 (green) and NHP 22117 (yellow)), supporting a model where Mtb dissemination is structured as parallel subnetworks of dissemination rather than a single radiation event. Further, we find 7 instances (Figure 3, arrows) where we can infer directional dissemination (i.e., where a mutation appears unfixed at an origin site and then hitchhikes as a fixed mutation to downstream tissues). Of these, 5 dissemination events involve Mtb spreading from and to lung sites (Figure 3, red arrows), while 2 cases involve dissemination to lung sites from a thoracic lymph node (Figure 3, blue arrows), suggesting that Mtb can in principle return to the lung after dissemination to other anatomical sites.

### Mtb mutations in NHPs and humans are enriched in lipid metabolism and biosynthetic processes

We next correlated the NHP intrahost mutations against two treatment-naïve human studies that also identified intrahost Mtb mutations: Liu et al.^17^ (321 mutations found in 257 genes in sputum from non-HIV infected patients with active TB); and Lieberman et al.^18^ (518 intrahost mutations in 403 genes from an autopsy study of HIV-coinfected individuals). In all 3 datasets, intrahost mutations were mostly found once in different genes, which precluded looking at the ratio of non-synonymous to synonymous mutations on a gene basis; and only the iron transporter, irtA, was mutated in all three datasets (Figure 4A). Despite lack of one-to-one gene overlap, there was a significant enrichment of intrahost mutations in Mycobroswer^26^-predicted lipid metabolism genes (Figure 4B) in the NHP dataset (15% of mutations) and Lieberman et al. (9% of mutations) datasets, with a statistical trend (p=0.05) for Liu et al. (10% of mutations).

**Figure 4.**
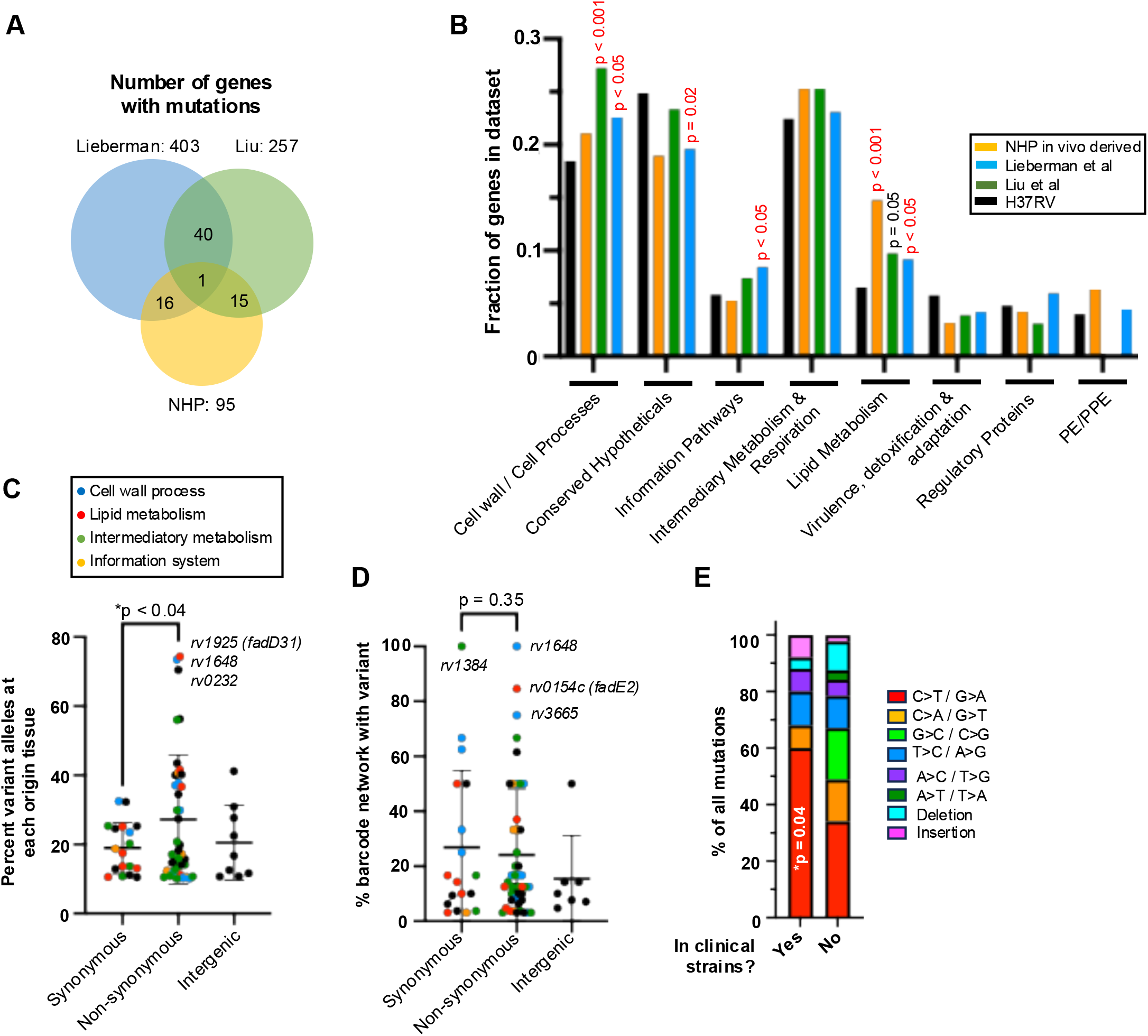
Intrahost mutations are enriched in lipid metabolism genes and confer Mtb fitness. (A) Overlap of genes with intrahost mutations (synonymous and non-synonymous) in NHPs (95 genes) and two human datasets (257 and 403 genes for Liu et al.^17^ and Lieberman et al.^18^). (B) Fraction of intrahost mutations in genes from each Mycobrowser category is plotted. The enrichment of intrahost mutations in each functional category was calculated against the number of genes annotated for each category in the H37Rv genome using a Fisher’s exact test. (C) The proportion of mutant reads versus all reads was compared at each origin tissue. (D) The percent of tissues containing a barcoded Mtb strain that also carried a genomic variant of interest was compared. (C-D) Statistical analyses were performed using an unpaired t-test. (E) A Fisher’s Exact test was used to compare the proportion of C-to-T transitions for variant alleles that were and were not found across 55,000 human clinical strain genomes.

Further, we find that intra-NHP mutated genes had significantly more predicted interactions than expected by chance (p = 0.005) using the STRING database^27^ and that the putative gene interactions clustered by MycoBrowser functional categories, with lipid metabolism genes (especially *pks* genes) centrally located (Supplementary Figure 2). InterPro analysis identified several protein domains that were enriched in the NHP dataset (Supplementary Figure 3), which included lipid metabolism domains (e.g., acyltransferase (FDR< 1x 10^-5^)) and polyketide synthase-associated domains (FDR < 3 x 10^-3^). The genes associated with these terms were highly overlapping and included *pks7, pks8, pks12, mbtD, pks1* and *pks2*, which are involved in the biosynthesis of diverse cell wall lipid structures and secondary metabolites. As in NHPs, we did observe enrichment of polyketide domains terms in the human datasets as well (Supplementary Figure 3), suggesting these biosynthetic genes as a whole are common targets of mutation in vivo.

### Intrahost mutations are enriched within granulomas and are associated with dissemination

We next assessed whether in vivo derived mutations can confer intrahost fitness benefits during infection, focusing on mutations that change protein sequence (missense, indels), which are more likely to impact gene function and Mtb fitness compared to synonymous mutations. To do this, we considered Mtb fitness across two anatomical scales: either locally within an individual tissue or systemically through dissemination to multiple tissues. First, to assess individual tissue fitness, we compared the proportion of wild type alleles to non-synonymous and synonymous mutations, finding that the prevalence of non-synonymous mutations were significantly higher than that of synonymous alleles (Figure 4C), suggesting that a subset of these mutations may be locally beneficial for Mtb. Second, to study dissemination, we compared the fraction of tissues infected by a barcoded Mtb strain that also contained a non-synonymous or synonymous genomic variant, with the hypothesis that dissemination-promoting mutations will be enriched within disseminated networks. Here, we did not observe a statistical difference between synonymous and non-synonymous mutations, though there is a wide range of dissemination in both groups (Figure 4D). Interestingly, mutations that were associated with highly disseminated Mtb strains are distinct from those that are enriched in individual tissues, which may suggest that additional factors beyond local fitness contribute to dissemination.

### NHP derived Mtb mutations also arise in human clinical strains

We finally assessed whether intrahost NHP mutations can be found across human Mtb clinical strains. To do this, we leveraged a recent phylogenetic analysis that mapped over 55,000 Mtb clinical genomes to an ancestral genome^28^ and looked for identical mutations as those observed in our NHP dataset. We find 22 mutations (12 non-synonymous, 9 synonymous and 3 intergenic) that were also present in at least one human Mtb clinical strain (Table 1). Most of these mutations appear in a single clinical isolate, but some mutations (e.g., pks7 frameshift at codon 1174 and non-synonymous mutations in *rv0785, rv0987* and *rv3594*) arose independently across different infections and were successfully transmitted between people (e.g., a missense mutation in *rnj*, a gene whose loss has been associated with drug tolerance^29^, arose once but was subsequently shared across 10 individuals). We also find an intergenic mutation 25 nucleotides upstream of the ESX-1 regulator *whiB6*, which is an area of high mutational diversity across clinical strains^28^. Finally, we show that intra-NHP mutations that match mutations in clinical strains are significantly enriched for C>T/G>A mutations (Figure 4E), further supporting a model in which immune oxidative damage is a driver of Mtb diversification in the host.

**Table 1.** NHP intrahost mutations are also observed in human Mtb clinical strains. Identical genomic mutations in the NHP dataset were matched to mutations observed in over 55,000 clinical isolates^28^. Clinical strain mutations and the number of independent mutations events were predicted by comparing each clinical strain against a Mtb ancestral genome reconstruction. Nonsyn = non-synonymous, Syn = synonymous, IG = intergenic. Pks7 mutations with an ‘A’ and ‘B’ indicate different mutations leading to a frameshift in the same codon, while we find two different types of intergenic whiB6 mutations at the same position.

| Gene | DNA change | Protein Change | Mutation | # Events | # Strains |
| --- | --- | --- | --- | --- | --- |
| rv0785 | C>CG | NonSyn | Ala561fs | 2 | 2 |
| rv0987 | A>G | NonSyn | Ser710fs | 3 | 3 |
| pks7 | CG>C | NonSyn | Ala1174fs | A: 5<br>B: 2 | A: 21<br>B: 3 |
| rv0075 | G>A | NonSyn | Gly175Ser | 1 | 1 |
| rv0943 | G>A | NonSyn | Ala54Val | 1 | 1 |
| rv1648 | A>G | NonSyn | Ser26Gly | 1 | 3 |
| pks8 | C>T | NonSyn | Thr1274Met | 1 | 1 |
| lipJ | G>A | NonSyn | Ala390Val | 1 | 4 |
| rv2075 | C>A | NonSyn | Val415Phe | 1 | 1 |
| rv2565 | C>T | NonSyn | Ser292Phe | 1 | 1 |
| rnj | T>C | NonSyn | Arg361Gly | 1 | 10 |
| entC | C>T | NonSyn | Ala270Val | 1 | 8 |
| rv3594 | G>A | NonSyn | Arg215His | 2 | 3 |
| fadD34 | C>T | Syn | Gly143Gly | 1 | 1 |
| Rv1520 | T>C | Syn | Arg81Arg | 1 | 1 |
| Rv1835c | C>T | Syn | Ala326Ala | 2 | 2 |
| Rv2402 | G>A | Syn | Ala324Ala | 1 | 1 |
| Rv2426c | T>G | Syn | Ala256Ala | 3 | 12 |
| Rv3228 | G>A | Syn | Leu102Leu | 2 | 6 |
| fadE31 | C>T | Syn | Asp190Asp | 2 | 4 |
| aftA | C>T | Syn | Arg5Arg | 5 | 27 |
| glfT2 | G>A | Syn | Asn229Asn | 1 | 3 |
| PE-PGRS38-pbpB | C>A | IG | PGRS38 (-20) | 1 | 1 |
| Rv3268-Rv3269 | A>C | IG | rv3269 (-110) | 1 | 2 |
| whiB6-Rv3863 | C>T | IG | whiB6 (-25) | C>T: 4<br>C>G:2 | C>T:4<br>C>G:10 |

## Discussion

Defining the intrahost Mtb mutational repertoire within treatment-naive hosts may inform the evolutionary pressures shaping Mtb adaptation to the host and also record how immune environments may differ in individuals who have other comorbidities such as HIV co-infection. Here, sequencing of Mtb from infected NHP hosts finds that SIV coinfection was associated with significantly increased Mtb mutational diversity compared to Mtb only animals. This appeared to be mainly driven by shorter generation times and greater overall Mtb replication capacity in SIV-infected hosts, which create more opportunities for mutations to occur. Conversely, increased oxidative immune pressure is not likely driving mutation, as chronic SIV coinfection is known to impair immune control in this NHP model^30,31^ and we observe fewer oxidative damage associated mutations in SIV animals relative to Mtb only controls, a finding that has also been reported in HIV-positive patients^17,18^.

Further, we also evaluated the impact of ART on Mtb evolution within SIV-infected hosts. Prior barcoding analyses suggested that SIV+ART+TB animals remained more permissive for Mtb dissemination to extrapulmonary sites, suggesting a level of continuing immune dysfunction despite virologic control^7^. Here, we find Mtb from a majority of ART animals had an increased proportion of oxidative damage associated mutations relative to TB only animals, and that this signal was positively correlated with higher overall mutational rates. We speculate that a subset of SIV+ART+TB animals could be mounting a dysregulated heightened inflammatory response to Mtb infection. Previous work has shown that alveolar macrophages from HIV-negative individuals on pre-exposure ART prophylaxis mount an altered immune response to subsequent Mtb exposure^32^ and as we did not have samples from an SIV-negative ART only group, we cannot discern if antiretroviral exposure alone or a combination of SIV and ART is responsible for this shift in Mtb oxidative damage. Beyond studying Mtb evolution during coinfection, we envision that structured population sequencing in combination with barcoding analyses can be leveraged in translational contexts such as in vaccine studies. For example, quantification of barcoded Mtb strains has found that intravenous BCG (IV-BCG) vaccination is associated with a long-lasting ability to restrict early Mtb bacterial establishment^33^, and it would be interesting to use Mtb population sequencing to assess whether bacteria in IV-BCG vaccinated hosts exhibit mutational profiles and trajectories reflective of enhanced immune pressure. Intrahost sequencing may also identify tissues enriched for Mtb mutation events that could be selected for deeper immune profiling and may aid discovery of novel immune correlates of protection.

Our findings also contextualize bacterial evolution and dissemination within the host. First, as granulomas are typically colonized by a single bacterial founder and that different Mtb isogenic strains have heterogeneous dissemination capacity even within the same host^34^, we were able to incorporate intrahost mutations as secondary ‘barcodes’ to further study otherwise clonally barcoded strains and infer complex dissemination patterns where Mtb is likely spreading in parallel subnetworks strains originating from different tissues. This is in contrast to a single event dissemination model, where all future disseminated sites are seeded from a single early lesion. We do note that a majority of the dissemination networks harbor mutations at a single tissue, which could be consistent with a single event model, but given that there are few intrahost mutations detected in general, we cannot rule out parallel dissemination in these cases as well.

Second, our genomic mapping found that intrahost mutations were enriched in Mtb lipid catabolism and biosynthetic genes (Figure 5), which may reflect the importance of beta-oxidation of host lipids for Mtb nutrient acquisition during infection^35^. While beta-oxidation supports carbon acquisition, it also increases reductive stress by generating NADH, which can be counteracted through several mechanisms. First, NADH is consumed through respiration, and we observe mutations in putative nitrite efflux proteins *narK3* and *narU*, which may facilitate the use of nitrate as a terminal electron acceptor in the context of microaerophilic conditions^36^. Second, NADPH equivalents are utilized for biosynthesis of cell envelope components and lipid storage molecules by large modular polyketide synthase (pks). The pks enzymes are responsible for producing key cell wall components (e.g., dimycocerosates, mycocerosic acids, lipoarabinomannan, mannosyl-ß-1 phosphomycoketides, sulfolipid and the siderophore mycobactin) using lipid precursors^37^; and recent work has shown that phthiocerol dimycocerosate production is required for optimal Mtb growth on propionate, which presumably acts to re-balance the reductive stress from odd-chain lipid degradation^38^. Here, we find significant enrichment of intrahost mutations—both in NHP and human datasets—within *pks* genes, which may be beneficial to Mtb in the human population; for example, we identified a frameshift in *pks7* that also appears to be independently mutated 7 times across human clinical strains and has been transmitted between 24 individuals (Table 1). Interestingly, while *pks7* knockout strains have been reported to be attenuated in aerosol infection in C57BL/6 mice^39^, our data would suggest this enzyme can be dispensable in human infection and transmission, which would be consistent with recent work showing *pks7* insertion mutants retaining the ability to intravenously infect a genetically diverse panel of mice^40^. Together, these mutational signatures highlight the metabolic flexibility of Mtb in balancing energy and redox needs under host-imposed stresses.

**Figure 5.**
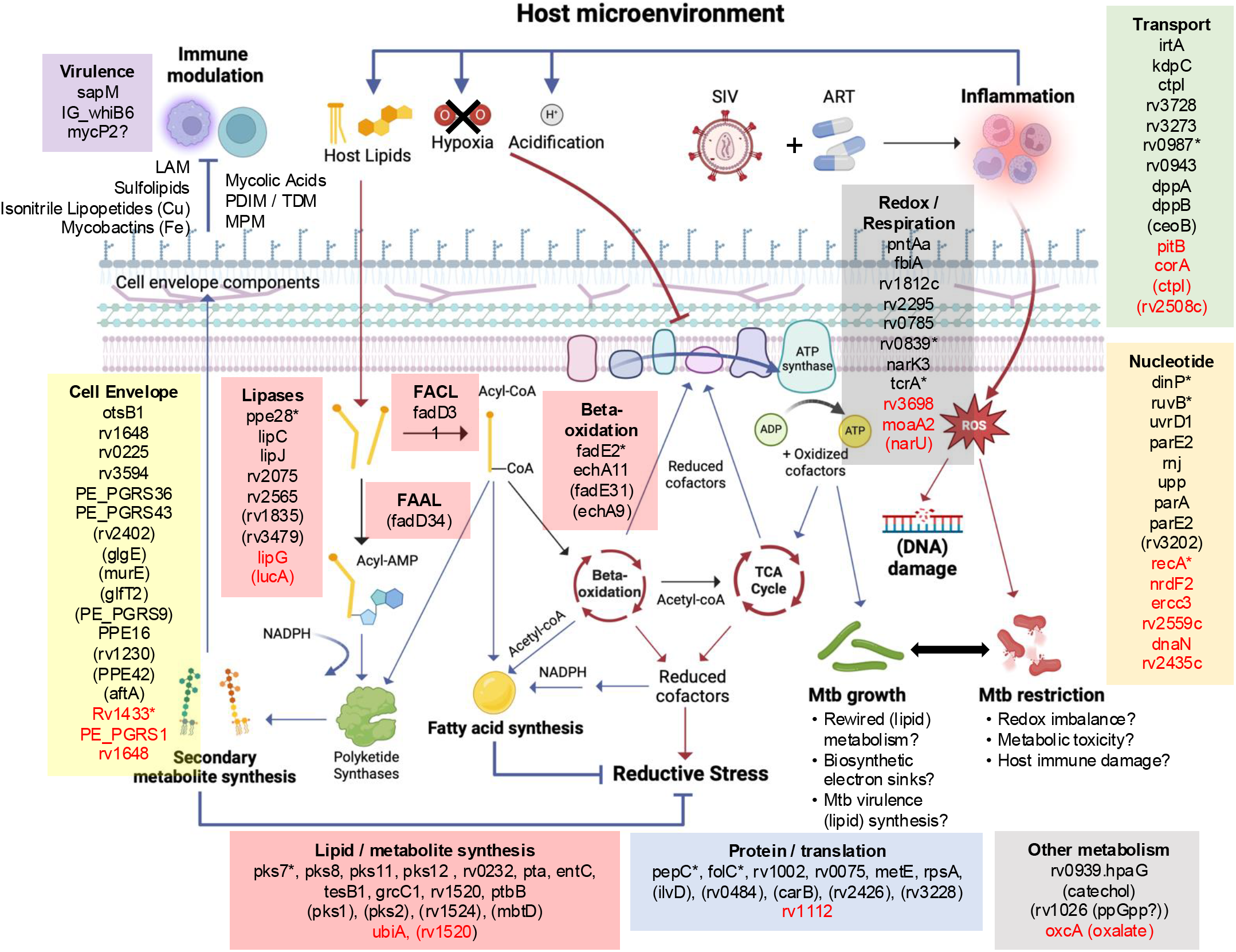
Model of intrahost mutations in metabolic and cell envelope processes. NHP-derived Mtb mutations are grouped into functional processes using literature curation and conserved domain similarity to microbial homologs. Genes in black represent all intrahost derived Mtb mutations, while genes in red are mutations assigned to the ‘pre-existing’ mutation group. Genes marked with asterisks represent frameshift mutations; genes in parenthesis represent synonymous mutations and the remaining are predicted missense mutations. Functional categories are colored as follows: lipid metabolism and biosynthesis (red), cell envelope (yellow), nucleotide synthesis/repair (orange), metabolite transport (green), protein homeostasis/translation (blue), redox processes/respiration (grey), virulence (purple). The schematic depicts lipases breaking down complex host and bacterial lipids into fatty acids, which can then be activated for degradation (or incorporated into other molecules) by conjugation to CoA through fatty acid CoA ligases (FACL). Different acyl-CoA substrates are then degraded through iterative cycles of beta-oxidation, yielding acetyl-CoA that can be used to drive the TCA cycle and respiration as well as being incorporated into Mtb secondary metabolites. Fatty acid ligation to AMP via fatty acid AMP ligases (FAAL) can be directly incorporated by pks (and other biosynthesis genes) into larger lipid-based structures, many of which are components of the cell envelope. Beta-oxidation of lipids also yields NADH, leading to increased reductive stress that needs to be re-balanced. This can occur through conversation of NADH to NADPH, which is utilized for secondary metabolite synthesis as well as NADH consumption during respiration, which can become impaired due to host pressures like hypoxia, acidification and inflammation. This may lead to leakage of electrons from the respiratory chain, generating reactive oxygen species and macromolecule damage in Mtb. In the context of ART treatment, we hypothesize an augmented inflammatory environment not seen in Mtb only animals, which may yield greater Mtb damage and increased detection of oxidative-damage associated mutations. Image was prepared using BioRender.

### Limitations to the study

Several limitations to this study remain. First, our dataset likely underestimates true intrahost mutational diversity, as late-emerging mutations (which will be at lower prevalence compared to early arising mutations) are unlikely to be sampled for the sequencing depth used in this study. Further, some variants will also have been filtered out by our 10% prevalence threshold, which was imposed to reduce false positive calling of low-frequency sequencing errors. Finally, highly repetitive genes of the PE/PPE families are difficult to uniquely map using short read sequencing technology, so we have masked these regions in our reference genome^20^ for subsequent analyses.

Second, our analyses are restricted to an early infection time point when the bacterial population size remains sizeable but mutational diversity may be limited compared to human infection with Mtb, where infections of months to years enable Mtb to accumulate intrahost mutations. On the opposite end, human longitudinal studies have shown that antibiotic treatment (and presumably immune attack leading to latent infection) will shrink the in vivo airway mutant pool^13–15^, which will likely eliminate subsets of mutations that were beneficial early—but not late—in infection. Further, this NHP study does not capture population bottlenecks occurring during inter-host transmission, where mutations that may be beneficial within a host also must successfully access the airway, survive exposure to the outside environment and transmit infection to a new host. This high bar for spread between hosts may manifest as strong transmission bottlenecks and explain why the vast majority of mutations found across over 55,000 Mtb clinical strains exist at terminal branches within single individuals^28^.

Finally, while gene-level data suggests ongoing Mtb mutation in lipid metabolic genes, it is difficult to infer whether a given missense or synonymous mutation is positively or negatively impacting gene function and thereby allow us to nominate pathways as essential or non-essential for host adaptation. Additionally, as the exact lipid substrates of many paralogues of genes involved in lipid metabolism and biosynthesis remain unknown, future work will be needed to define the nutrient diversity of Mtb host microenvironments—especially at relatively inaccessible anatomic sites beyond sputum—that promote and restrict successful Mtb infection.

## METHODS

### Non-human primate study approval

This is a secondary use study employing existing genomic DNA samples from a prior study that involved non-human primates^7^. All animal procedures were approved by the University of Pittsburgh IACUC in compliance with the Animal Welfare Act and Guide for the Care and Use of the Laboratory Animals.

### Sequencing of Mtb genomic populations from infected NHP tissues

We curated a set of Mtb genomic DNA samples that were previously generated for barcode sequencing^7^. In order to compare Mtb evolution across similar infection time scales, we chose to analyze samples only from the 20 NHPs from the original study that survived 10-12 weeks of Mtb infection, though we did sequence tissues from 3 additional SIV coinfected NHPs that reached a humane endpoint much earlier. Whole genome sequencing was performed on the Illumina NovaSeq S4 300 cycle platform across 4 sequencing runs (see Supplementary Table 2 for metrics and sequencing accession codes). Reads were filtered for good sequencing quality reads using fastp^41^ and then mapped to the recently generated barcoded Mtb Erdman reference genome (GenBank Accession CP172229.1)^20^, which was obtained from an aliquot of the library used to infect NHPs across studies. After alignment, we applied Mutect2 from the GATK toolkit (https://gatk.broadinstitute.org/hc/en-us) to identify all mutational changes (single nucleotide polymorphisms and insertion/deletions) at any frequency relative to the reference genome. The functional effect of each mutation (e.g. intergenic, synonymous, missense, frameshift) was also predicted using SnpEff based on our recent Erdman annotation^20^. We included a mask to redact variants predicted in repetitive sites in our Erdman reference genome that had been predicted to have low mappability^20^, as they are more likely to arise from mismapping events.

The Mutect2 output yielded a preliminary list of over 100,000 detected genomic changes across all samples, which were mainly found at low prevalence (<10% mutant alleles versus wildtype sequences) and driven primary by a subset of samples. Due to the logistics of tissue sampling, each infected tissue was homogenized for Mtb plating and the entire Mtb population was collected for gDNA purification. Therefore, we lack technical replicates from the same sample to assess the impact of tissue homogenization and bacterial gDNA extraction on introducing DNA mutations. However, we aimed to filter out the vast majority of potential artefactual mutations (though false positives may still remain), by enacting 3 QC steps: first, we used the interquartile range (IQR) method to define outlier samples with a number of Mutect2 variants at 1.5x greater than the IQR across all the samples from each sequencing batch. This yielded 46 outlier samples that were removed from subsequent analyses (Supplemental Figure 1A, ‘QC1’). Notably, the vast majority of mutations in these outliers were enriched for C to A transversions (Supplemental Figure 1B), a mutational change that has been previously associated with DNA damage artifacts arising during library preparation due to elevated heat^23^. Second, we filtered remaining variants to remove low-confidence calls at sites with substandard sequencing depth, which included only included 1) sequencing runs with an average read depth above 50x; 2) genomic variants that sat in regions where there was overall sequencing depth that exceeded 50% of the average sequencing depth of the sample; and 3) variants whose prevalence was above 10% (relative to all reads at that position) to account for rare sequencing errors (Supplemental Figure 1A, ‘QC2’). Given the stochastic nature of oxidative damage during library preparation, which should yield rare mutations, we also expect the minimum prevalence threshold will aid in removal of library preparation artefacts. Third, due to difficulty in uniquely assigning short reads to repetitive regions in the Mtb genome, we reasoned that there will be systematic mismapping events yielding the same exact ‘mutation’ (in both chromosomal position and type of DNA base change) being found in samples even across different animals. Filtering out the exact same variants found in more than 1 animal yielded a candidate list of 217 variants found across 125 tissues (Supplemental Figure 1A, ‘QC3 step’). Finally, we visually inspected the read pileups generated by each mapping file using Samtools and confirmed that the different reads were mapping across the site of interest and that variants were not clustering at the 5’ and 3’ ends of reads. This produced our final validated table of 143 unique high confidence de novo mutations for further analysis (Supplementary Table 2).

### Defining pre-existing and intrahost-derived Mtb mutations

As our reference Erdman genome was derived from a single barcoded clone within a larger barcoded library, we cannot exclude that some mutations identified by sequencing could have arisen in vitro and is specific to a given barcoded strain. To conservatively separate in vivo derived mutations versus those that may have been pre-existing in the library, we compiled the Mtb barcoded strains present in each tissue sample from prior barcode amplicon sequencing data^7^ and compared this to the Mutect2 detected variants present in each tissue. As diagramed in Figure 1B, we considered mutations in vivo-derived if they satisfied 1 of the 2 following criteria.

First, we first looked for tissues that contained a genomic variant and a single barcoded Mtb strain, which allowed us to link a genomic variant to a barcoded Mtb strain genome. We then identified all tissues containing that barcoded Mtb strain and assessed the presence/absence of the genomic variant of interest in those tissues. If we observed a pattern where some tissues had only the wild type allele while others had the genomic variant, we reasoned the detected mutation likely originated during infection. This situation is diagrammed as ‘BC2’ in Figure 1B.

Second, when we cannot link a genomic variant to a single barcoded Mtb strain (often the case when a mutation is found in a tissue that was infected by more than one barcoded Mtb strain), we looked for the genomic variant of interest across all tissues containing every possible barcoded Mtb strain it could be linked to. If we observed both presence of wild type Mtb and mutant alleles in every potential barcode network, we also considered this variant as likely emerging in vivo. This situation is diagramed as ‘BC3’ and ‘BC4’ in Figure 1B.

If a variant does not satisfy the criteria above, we cannot rule out the mutation could have emerged in vitro, so it is considered ‘pre-existing’ and excluded from further analyses. An example of this is ‘BC1’ in Figure 1B, where a mutation was present in all tissues that was infected that barcoded strain.

### Defining mutant origin tissues and timing of origin emergence

To define mutation ‘origin’ sites, we first identified tissues containing a single barcoded Mtb strain. If sites contain an unfixed mutant allele (i.e., <75% of all reads at that site are mutant), then we considered these tissues as harboring both wildtype (i.e., parental) and mutant alleles. We elected to use a 75% threshold based on the WHO threshold used to define high-confidence mutations associated with drug resistance^42^, and empirically found that using a 90% threshold would also not change the origin site designations in this study. These are likely the origin sites where the mutation originally arose. In the parent study, animal lungs were imaged using PET-CT on a monthly basis at 4-, 8, and 12-weeks post infection to register the location and presence of newly detected lung lesions^7^. Here, we compared the time of PET-CT detection of the lung-specific origin sites (PET-CT registration was not done for thoracic lymph node and extrapulmonary sites) across treatment groups.

### Bacterial burden quantification and flow cytometry analysis of lesions across animals

We obtained previously published tissue-level live bacterial burden (CFU) and total (live and dead) bacterial chromosomal equivalents (CEQ) measurements for the animals that were included in this study^7^. Total CFU and CEQs were summed for all sampled tissues from each animal for statistical comparisons. We also obtained lesion-level flow cytometry profiling data from the prior study^7^, in which CD11b-positive myeloid cells were profiled for expression of activation and cytokine markers.

### Mutation rate quantification

Similar to previous work using colony based sequencing from infected NHPs^16^, we calculated a Mtb mutation rate for each animal by dividing the number of unique mutational events (after normalizing for number of tissues sampled) by the number of bacterial generations (per unit of infection time) occurring in the host. Here, we summed the number of different mutations across all tissues in a NHP to estimate the mutation rate of the entire Mtb population within one animal using the following equation: ***μ* = *m* /⍰[*N*⍰*⍰(*t*⍰⍰⍰*g*)] *D***

In this equation, <u>μ</u> represents the total mutation rate across an animal, <u>m</u> is the number of unique mutational events detected by sequencing across all tissues in the animal, <u>N</u> is the Mtb genome size (set at 4,400,000 bp), *t* is the total infection time of each NHP (days of infection multiplied by 24 hours), <u>g</u> is the Mtb in vivo generation time (in hours) and <u>D</u> is the total number of tissues that were sequenced per animal. We applied two versions of this calculation. First, to establish a relative mutation frequency per day of infection (Figure 2A), we used a fixed 18 hours for a single doubling across all treatment groups. The numerical values calculated with a fixed doubling time is the <u>*relative*</u> mutation frequencies per fixed unit of infection time, which was used to establish a fold change difference between treatment conditions (relative to the mean mutation frequency in the Mtb only group). Second, to calculate a specific Mtb mutation rate per generation (Figure 2C), we first estimated the average Mtb generation time (g) in each NHP using the following formula: **g = t (log2 (final Mtb population size founding Mtb population size))**. In this equation, we used the total number of Mtb CEQs obtained across all infected tissues in each animal as the final Mtb population size and the number of unique barcodes found in each NHP as the starting Mtb founder population size. We then used a NHP-specific generation time (g) to calculate the Mtb mutation rate for each animal using the first equation.

### Bacterial barcoding network visualization

Using previously published amplicon barcoding data^7^ from the tissues sequenced in this study, we generated network plots by converting a table of genomic variants present in all sequenced tissues into a network plot using the R package, igraph. Specifically, each tissue that contained a given barcoded Mtb strain was initially plotted as nodes given the same color. A ring was added in a different color to a tissue/node if it contained a unique genomic variant after sequencing such that different variants within the same barcoded dissemination network are represented by different colored rings. Using the presence or absence of genomic mutations shared across tissues, we inferred dissemination subnetworks with the fewest dissemination events. In the case of networks for which we are unable to order the events of dissemination, we arbitrarily represented dissemination as a single star-like network. If a genomic variant was found in more than one tissue with a network, we manually re-ordered connecting lines between these nodes to form sub-networks. We finally imported the igraph barcode networks into Adobe Illustrator to visually improve line definition, update colors for visualization and scale for size. The edgelist representing the connections between tissues for all visualized barcoded networks is provided as an R object in the data folder on the Github page: https://github.com/Fortune-Lab/Mtb-NHP-genotyping. Note: some tissues harbored multiple genomic mutations but igraph is limited to adding a single ring, so we added these variants and the site of origin sites (as boxes) in Adobe Illustrator. The presence of genomic mutations and origin sites within the networks are indicated by metadata columns in the R object.

### STRING and Interpro analyses

The list of all genes with in vivo derived mutations from our NHP data, Liu et al.^17^ and Liberman et al.^18^ were analyzed on the STRING database (https://string-db.org) to define putative interactions. All interacting nodes were then colored by their predicted^26^ functional categories. Finally, protein domain enrichment analysis (InterPro) was performed on the STRING database for all genes.

### Identification of intra-NHP mutations in human Mtb clinical strains

We obtained the list of genomic mutations identified across approximately 55,000 publicly available Mtb clinical isolate sequences from a recent publication^28^, which mapped mutations against a reconstructed ancestral Mtb genome and constructed a phylogenetic tree to identify independent mutational events and sharing of mutations across individuals. We searched for NHP derived intragenic mutations that also showed up with the same DNA mutation (e.g. C>T) at the same codon position in the human Mtb clinical isolate data. For intergenic sequences, we used BLAST to search for identical stretches of genomic sequence around the mutation site and then identified the relevant genomic position of the mutation in the ancestral Mtb genome used to map the human clinical isolates.

## Data availability

The genomic mapping, quality control and variant detection code and methods are fully described at the following Github link: https://github.com/Fortune-Lab/Mtb-NHP-genotyping. All sequencing data has been deposited on the Sequencing Read Archive, with accession codes and links for individual samples provided in Supplementary Table 1. All sequencing data are also available under Bioproject Accession PRJNA1432747. All metadata associated with the datasets in this work are deposited on Fairdomhub at the following link: https://fairdomhub.org/studies/1401

## Acknowledgments

This project has been funded in part with federal funds from the National Institute of Allergy & Infectious Diseases, National Institutes of Health under contract 75N93019C00071 and NIH, R01AI134195. The funders had no role in study design, data collection and interpretation, or the decision to submit the work for publication.

## FIGURE LEGENDS

**Supplementary Figure 1**. (A) The number of genomic variants (parentheses) detected by Mutect2 per sequenced sample (dots) before and after quality control (QC) steps. (B) The time of PET-CT detection (either 4-, 8- or 12-weeks post infection) of a lung granuloma that served as an origin site for an in vivo mutation is compared across groups. Statistical test compared the distribution of 8 and 12-week lesions using a Kruskal-Wallis test with Dunn’s correction. (C) STRING analyses showing the predicted interactions between all genes in the NHP dataset. Only interacting nodes are shown and are colored by their Mycobrowser category.

**Supplementary Figure 2**. All genes with intrahost mutations (all mutations = synonymous and non-synonymous mutations) for each dataset was analyzed for InterPro protein domain enrichment on the STRING database.

**Supplementary Figure 3**. All genes with intrahost mutations (all mutations = synonymous and non-synonymous mutations) for each dataset was analyzed for InterPro protein domain enrichment on the STRING database.

**Supplementary Table 1**. List of genomic accession numbers for all sequenced Mtb populations from infected tissues in this study. All samples are also available under the Bioproject accession: PRJNA1432747.

**Supplementary Table 2**. Final table of high confidence Mutect2-detected genomic mutations occurring in vivo.

