## Supplementary Figure 1 for "Population-level genome sequencing reveals distinct Mycobacterium tuberculosis intrahost mutational trajectories in simian immunodeficiency virus co-infected and antiretroviral treated non-human primates"

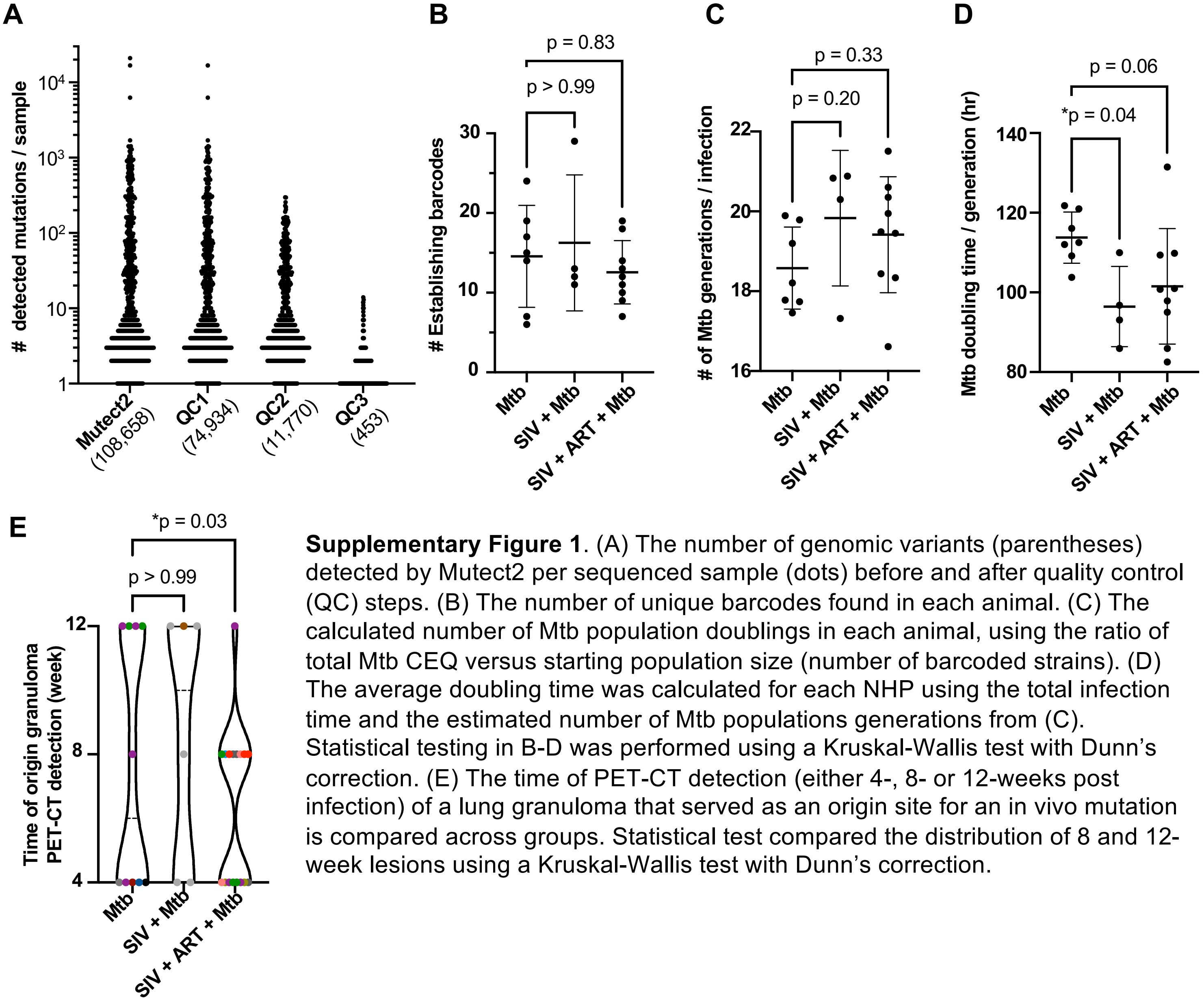


**Supplementary Figure 1**. (A) The number of genomic variants (parentheses) detected by Mutect2 per sequenced sample (dots) before and after quality control (QC) steps. (B) The number of unique barcodes found in each animal. (C) The calculated number of Mtb population doublings in each animal, using the ratio of total Mtb CEQ versus starting population size (number of barcoded strains). (D) The average doubling time was calculated for each NHP using the total infection time and the estimated number of Mtb populations generations from (C). Statistical testing in B-D was performed using a Kruskal-Wallis test with Dunn's correction. (E) The time of PET-CT detection (either 4-, 8- or 12-weeks post infection) of a lung granuloma that served as an origin site for an in vivo mutation is compared across groups. Statistical test compared the distribution of 8 and 12-week lesions using a Kruskal-Wallis test with Dunn's correction.
