## Supplementary Figure 2 for "Population-level genome sequencing reveals distinct Mycobacterium tuberculosis intrahost mutational trajectories in simian immunodeficiency virus co-infected and antiretroviral treated non-human primates"

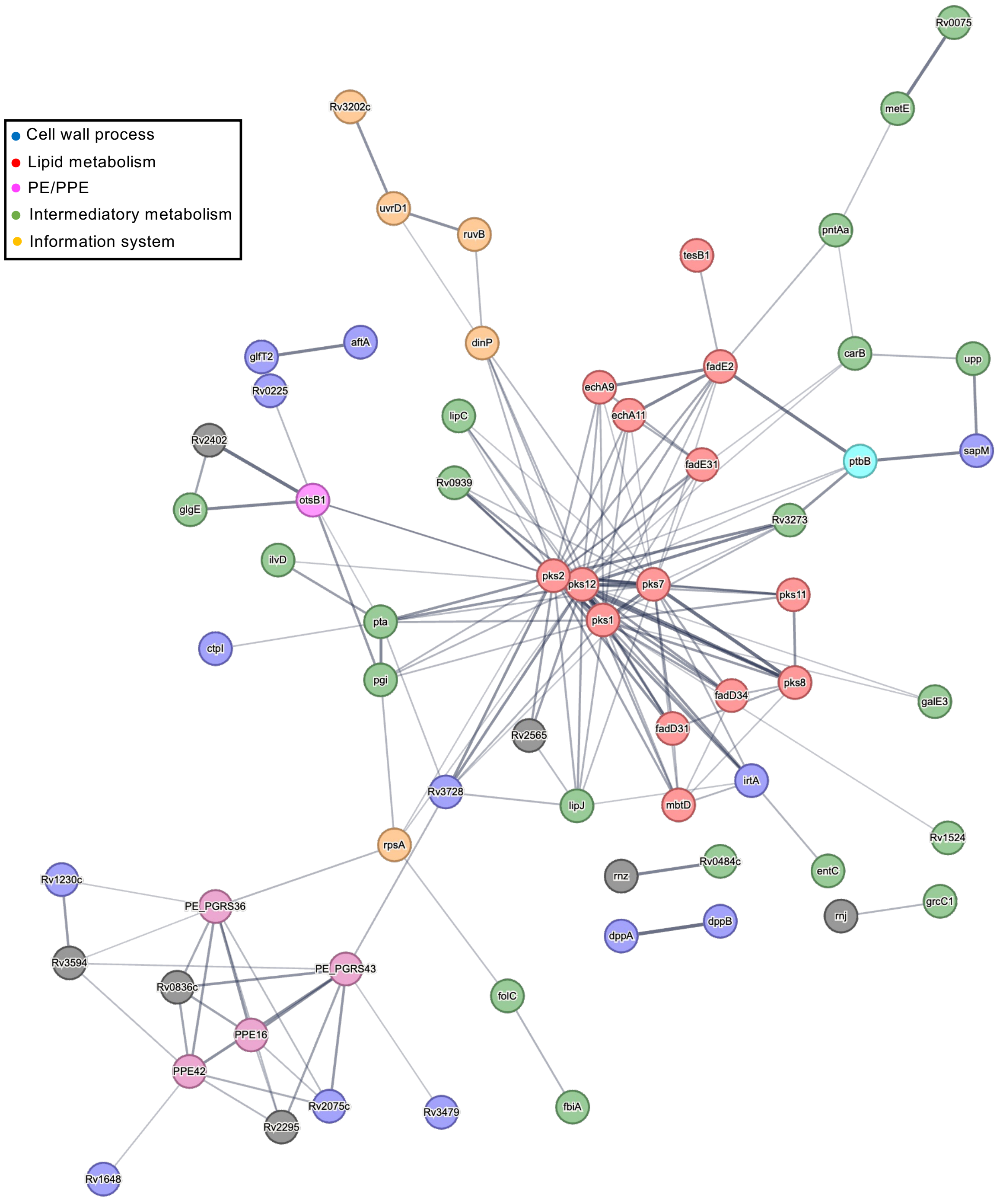


**Supplementary Figure 2**. STRING analyses showing the predicted interactions between all genes in the NHP dataset. Only interacting nodes are shown and are colored by their Mycobrowser category. All genes with intrahost mutations (all mutations = synonymous and non-synonymous mutations) for each dataset was analyzed for InterPro protein domain enrichment on the STRING database.
