## Supplementary Figure 3 for "Population-level genome sequencing reveals distinct Mycobacterium tuberculosis intrahost mutational trajectories in simian immunodeficiency virus co-infected and antiretroviral treated non-human primates"

**Supplementary Figure 3.** All genes with intrahost mutations (all mutations = synonymous and non-synonymous mutations) for each dataset was analyzed for InterPro protein domain enrichment on the STRING database.

### NHPs (all mutations)

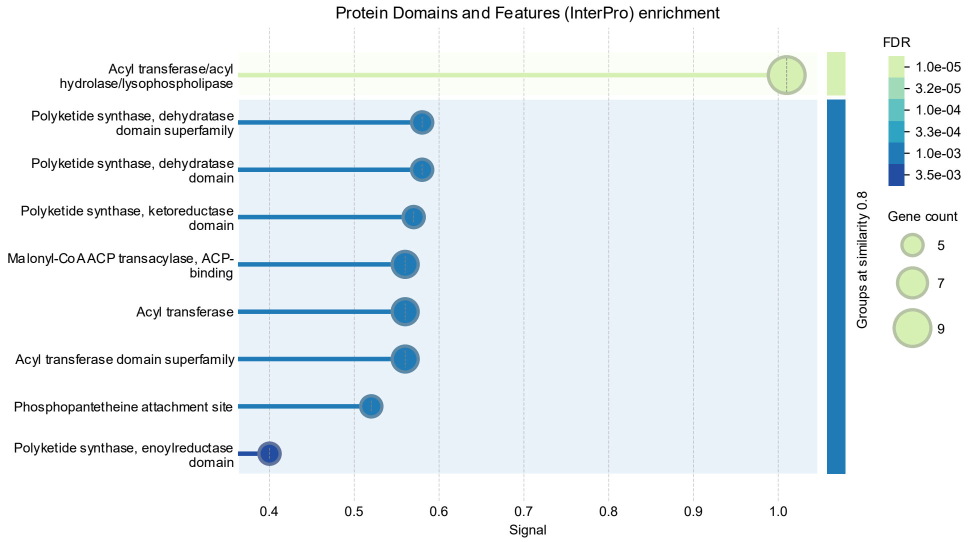


### Liu et al. (all mutations)

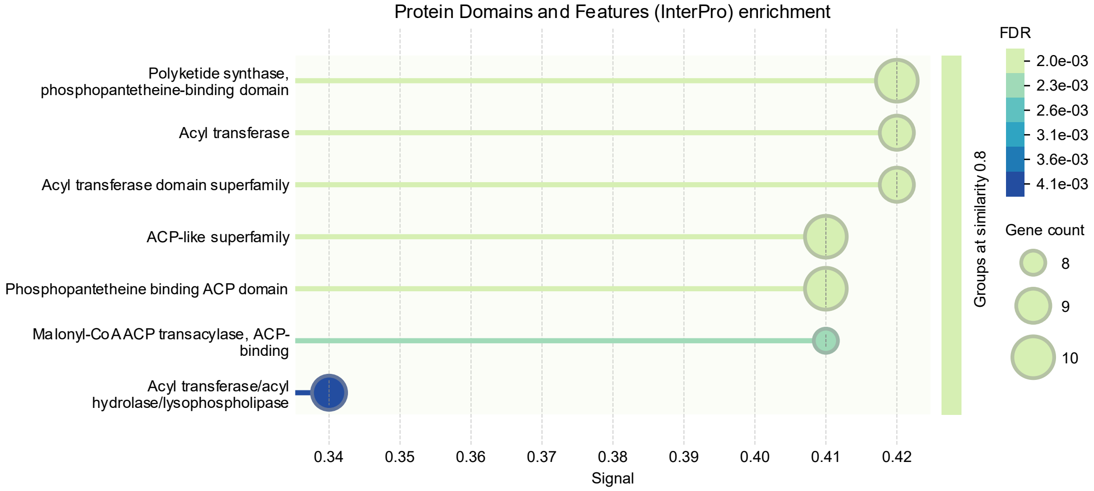


### Lieberman et al. (non-synonymous mutations)

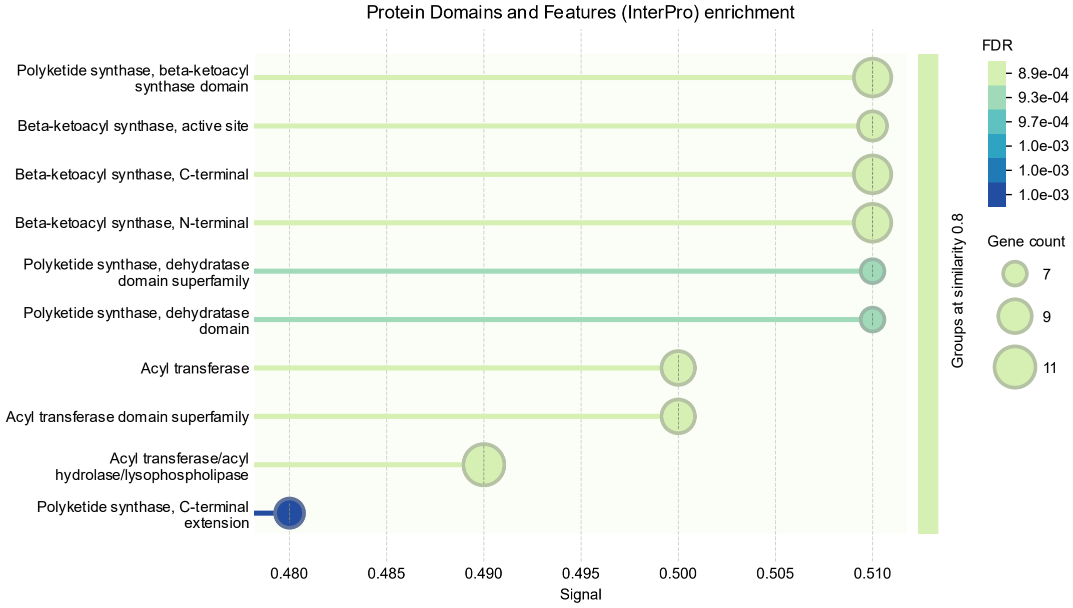
